# The influence of visual experience and cognitive goals on spatial representations of nociceptive stimuli

**DOI:** 10.1101/623561

**Authors:** Camille Vanderclausen, Louise Manfron, Anne De Volder, Valéry Legrain

**Affiliations:** Institute of Neuroscience, Université catholique de Louvain, Brussels, Belgium; Psychological Sciences Research Institute, Université catholique de Louvain, Louvain-la-Neuve, Belgium

## Abstract

Localizing pain is an important process as it allows detecting which part of the body is being hurt and identifying in its surrounding which stimulus is producing the damage. Nociceptive inputs should therefore be mapped according to both somatotopic (“which limb is stimulated?”) and spatiotopic representations (“where is the stimulated limb?”). Since the limbs constantly move in space, the brain has to realign the different spatial representations, for instance when the hands are crossed and the left/right hand is in the right/left part of space, in order to adequately guide actions towards the threatening object. Such ability is thought to be dependent on past sensory experience and contextual factors. This was tested by comparing performances of early blind and normally sighted participants during nociceptive temporal order judgment tasks. The instructions prioritized either anatomy (left/right hands) or the external space (left/right hemispaces). As compared to an uncrossed hands posture, sighted participants’ performances were decreased when the hands were crossed, whatever the instructions. Early blind participants’ performances were affected by crossing the hands only during spatial instruction, but not during anatomical instruction. These results indicate that nociceptive stimuli are automatically coded according to both somatotopic and spatiotopic representations, but the integration of the different spatial reference frames would depend on early visual experience and ongoing cognitive goals, illustrating the plasticity and the flexibility of the nociceptive system.

## 1. Introduction

One of the most relevant yet unresolved questions in pain research is how the brain localizes pain on the body [38]. Most research so far focused on the ability to localize nociceptive stimuli based on the somatotopic organization of the brain [1; 10; 11; 13; 32], anatomically mapping the body surface from the ordered projections of receptive fields [42]. However, as the body limbs are constantly moving in external space, such a spatial representation might be inefficient to appropriately localize the harmful stimulus around the body [38]. The spatiotopic representation considers the relative position of the body part receiving the stimulus in external space [52]. Using external space as reference frame, it allows the brain to identify the object in contact with the body, and spatially guide actions towards this object [14], such as defensive behaviors [29; 38].

The existence of the spatiotopic representation was assessed, mainly for touch and lately for nociception, using temporal order judgment (TOJ) tasks [7; 9; 18; 20–22; 30; 44–46; 54], during which participants determined the order of appearance of two successive somatosensory stimuli, one applied to each hand. TOJ are typically less accurate when their hands are crossed, as compared to an uncrossed posture, when reporting which hand is perceived as having been stimulated first. That effect is accounted for by the fact that the somatotopic representation mismatches the spatiotopic one (when crossed, a contact on the left hand is coming from the right part of space) [54]. The sensitivity of TOJ to posture indicates that nociception and touch, in addition to the somatotopic coding, are automatically coded according to spatiotopic coordinates taking into account the location of the hands in external space [30]. Additionally, crossing the hands decreases the perceived intensity of nociceptive stimuli [27; 51], highlighting the crucial role of spatial representations in pain processing. Interestingly, this automatic and default spatiotopic coding of somatic stimuli is not innate but shaped by early visual experience [30]. Accordingly, tactile TOJ performance of congenitally blind individuals is not affected by crossing the hands [18; 20; 44], witnessing their preferential reliance on somatotopic representations to localize somatosensory inputs [19].

Using TOJ tasks, we investigated the role of (1) past visual experience – by comparing performances of sighted and congenitally blind participants – and (2) ongoing cognitive goals – by manipulating spatial coordinates’ priorities through task instructions – in shaping the spatial representations of nociceptive stimuli. In addition to anatomical instruction, participants responded according to the side of space of the stimulus (spatial instruction). An absence of instruction effect in sighted participants would suggest that somatotopic and spatiotopic representations are equally competitive [9]. Alternatively, an improvement of TOJ performances in crossed posture during spatial instruction would suggest that the spatiotopic representation overrules the somatotopic one [54]. If early visual deprivation influences the development of the spatiotopic reference frame, spatial instruction should give rise to a crossing hand effect in blind participants. The absence of instruction effect would conversely suggest more flexibility in early blind people in shifting between the different spatial representations of nociceptive inputs according to cognitive goals [21].

## 2. Methods

### 2.1. Participants

Sixteen healthy sighted participants took part in Experiment 1. One participant was excluded because she could not properly achieve task requirements (see Analyses). The remaining fifteen participants (12 women) had a mean age of 24 years old (*SD*=2.6). Twelve of these participants were right-handed, according to the Flinders Handedness Survey [41]. Ten early blind participants (mean age=38, SD=13; 1 woman; 8 right-handed, 1 ambidextrous) took part in Experiment 2. Ten normally sighted participants were recruited as controls and matched to blind participants according to age and gender (Mean age=37, SD=13; 1 woman, 8 right-handed). None of the participants reported prior history of severe neurological, psychiatric or chronic pain disorders, traumatic injury of the upper limbs in the last six months, cutaneous lesion of the hands’ dorsa, regular use of psychotropic drugs, as well as intake of analgesic drugs (e.g. NSAIDS and paracetamol) within the twelve hours preceding the experiment. Normally sighted participants had normal or corrected vision. Early blind participants were recruited according to blindness attributed to peripheral deficits (see Table 1 for a complete description of blind participants). They were all considered as totally blind from birth. One of them (participant EB6) had very poor vision from birth and became definitively and totally blind consecutive to enucleation of the eyes at 18 months; he was therefore considered as early blind. Written informed consents were obtained for all participants and all experimental procedures were approved by the local biomedical ethics committee conformed to the Declaration of Helsinki. All participants received financial compensation for their participation.

**Table 1.** Description of the early blind participants.

| Participant | Gender;<br>Age | Handedness | Profession/<br>Educational<br>level | Visual<br>perception | Onset of<br>blindness | Cause of blindness | Braille | Cane | Musical<br>instrument | Musical<br>experience | Other |
| --- | --- | --- | --- | --- | --- | --- | --- | --- | --- | --- | --- |
| EB1 | M;29 | R | College degree | None | Birth | Leber congenital amaurosis | Yes | Yes | No | / | Various sport activities |
| EB2 | M;23 | R | Some college | None | Birth | Congenital cause (*) | Yes | Yes | No | / | / |
| EB3 | M;41 | R | College degree | None | Birth | Leber congenital amaurosis | Yes | Yes | Piano/organ | >5 years | Torball |
| EB4 | M;41 | R | Teacher/<br>College degree | None | Birth | Anterior chamber cleavage syndrome (Peters syndrome) | Yes | Yes + Dog | Piano/Drums/Flute | >5 years | Hearing aids |
| EB5 | M;32 | L | Some college | None | Birth | Hereditary retinal dysplasia | Yes | Yes | Guitare | >4 years | Slight auditory loss |
| EB6 | M;66 | R | College degree | None | 18 months | Bilateral retinoblastoma | Yes | Yes | Organ | >9 years | / |
| EB7 | F;36 | R | Phone sale until 2006/<br>Official high school | None | Birth | Severe corneal dysplasia | Few | Yes + Dog | Drums | Only few months | / |
| EB8 | M;33 | R | Some college | None | Birth | Persistent hyperplastic primary vitreous involving both eyes | Yes | Yes | Clavier | >10 years | / |
| EB9 | M;50 | R | College degree | None | Birth | Leber congenital amaurosis | Yes | Yes | Piano | >3 years | / |
| EB10 | M;26 | Ambi | Receptionist / Official high school | None | Birth | Leber congenital amaurosis | Yes | Yes | Drums, piano, harmonica, guitar | >5 years | ADHD syndrome |

EB: early blind; Age in years; F= female; M= male; R= right; L= left; Ambi=ambidextrous; (\*): no additional details available.

### 2.2. Stimuli and apparatus

Nociceptive stimulations consisted of radiant heat stimuli delivered onto the skin of the hands’ dorsa by means of two infrared CO_2_ laser stimulators (wavelength 10.6 *µ*m) (Laser Stimulation Device; SIFEC, Ferrières, Belgium). The power of the output stimulation was regulated using a feedback control based on an online measurement of the skin temperature at the site of stimulation by means of a radiometer whose field of view was collinear with the laser beam (see [16]). This allows defining specific skin temperature profiles. The laser beams were conducted through 10-m optical fibers. Each fiber ended with a head containing the optics used to collimate the laser beam to 6 mm diameter at the target site. Each laser head was hold upon each participant’s hand by means of articulated arms attached to a camera tripod system (Manfrotto, Cassola, Italy). Each laser head was fixed into a clamp attached to a 3-way head affording displacements of the target site of the laser beam perpendicularly oriented to the hand’s dorsum by means of several sliders going in all directions. Laser beams were displaced after each stimulus. Stimuli duration was 100ms. Stimuli were composed of a 10-ms heating ramp dedicated to reach the target temperature, followed by a 90-ms plateau during which the skin temperature was maintained at the target temperature. Heating was then stopped. The target temperature was determined for each participant’s hand according to individual activation threshold of nociceptive thinly myelinated Aδ fibers. Thresholds were estimated by means of an adaptive staircase procedure using reaction times (RTs) to discriminate detections triggered by Aδ-fiber inputs (RT < 650 ms) from detections triggered by C-fiber inputs (RT ≥ 650 ms) [16]. Participants were asked to press a button with the non-stimulated hand as soon as they felt something on the stimulated hand. Any RT equal or superior to 650ms leaded to a temperature increase of 1°C for the next stimulus. On the contrary, any RT inferior to 650ms leaded to a decrease in temperature of 1°C. The procedure started at 46°C and lasted until four reversals were encountered. The mean value of the four temperatures that leaded to a reversal was considered as the threshold. For the stimuli used during the experimental phase, 5°C were added to that threshold value, and if necessary, the temperatures were slightly adapted for each hand so that stimuli were perceived as equally intense between both hands. For sighted participants, view of the hands was prevented during threshold estimation. Stimuli at such temperature values were perceived as pricking and elicited a slightly painful sensation. Before each block of stimuli, sensations were tracked using a list of words to be chosen in order to describe the sensation (*not perceived*, *light touch*, *tingling*, *pricking*, *warm*, *burning*), and subjective intensity was rated using a numerical scale (from 0 (*no sensation*) to 10 (*strongest sensation imaginable*)). This was made to ensure that stimuli were still perceived as pricking and equally intense between the two hands. Stimuli temperatures were then adapted if necessary.

### 2.3. Procedure

The same procedure was used for Experiments 1 and 2. Participants were sitting on a chair, hands’ palms laying down on a table in front of them. During the uncrossed hands posture condition (see below), the tips of the two index fingers were separated by a distance of ~30cm. During the crossed hands posture condition, the same distance separated the tips of the two forth fingers. Distance between the reference fingers and the edge of the table was of ~40cm (Figure 1). Participants’ head was placed in a chin-rest in order to minimize head movement during the experiment. Noises from experimental devices were covered by a white noise played through earphones that the participants wore during the whole experiment. The sighted participants were blindfolded with an eye mask.

**Figure 1.**
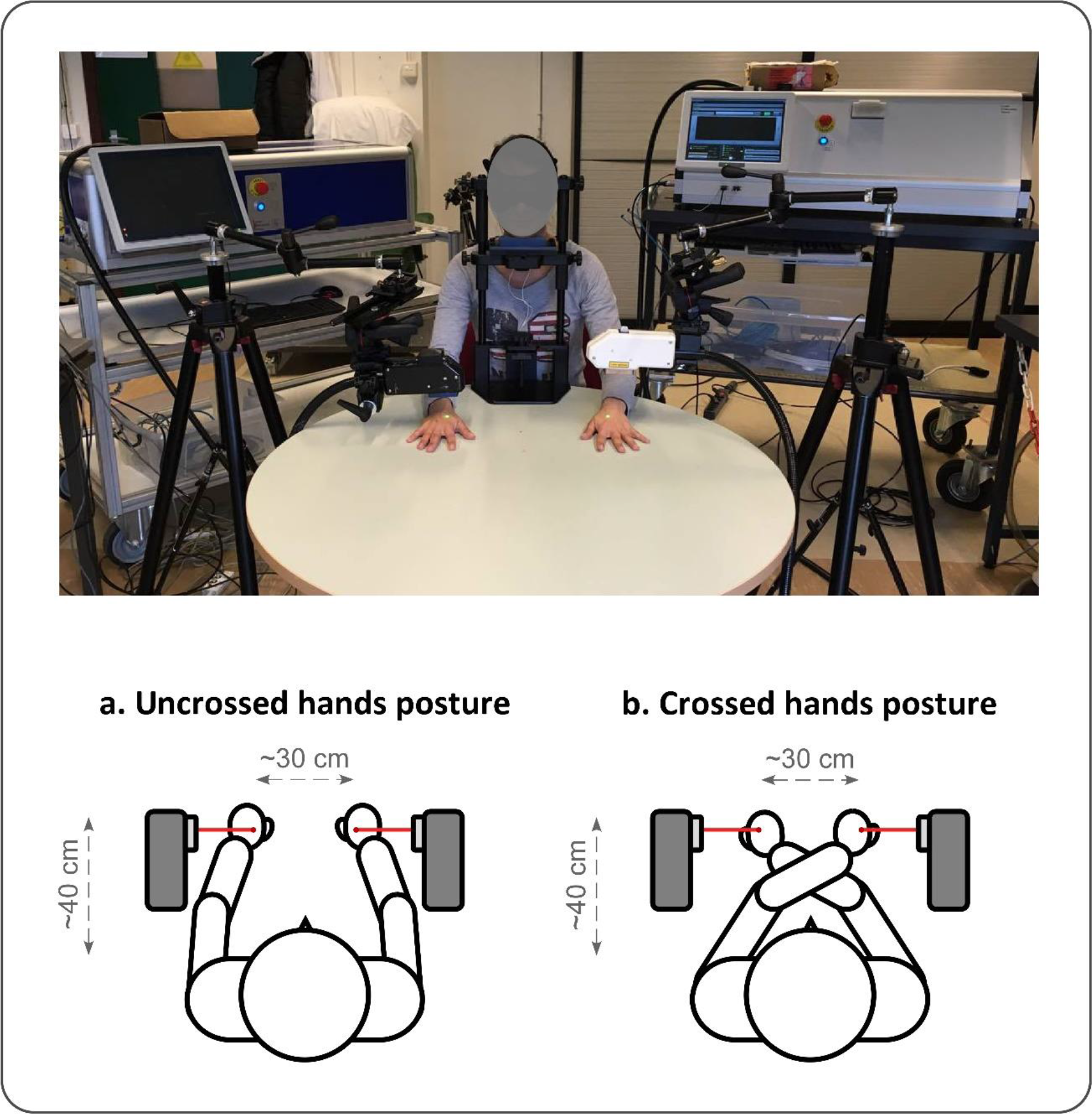
Experimental design of Experiments 1 and 2. Two temperature-controlled CO_2_ laser stimulators were used to activate Aδ-fibers. Laser beams were displaced after each trials on the hands’ dorsa using camera tripod systems. Temporal order judgement tasks were performed with the hands in either an uncrossed (A) or a crossed (B) posture. Sighted participants performed the tasks blindfolded.

Participants were presented with four blocks of 40 trials each. Each trial consisted in pairs of nociceptive stimuli, one applied on each hand, separated by 24 possible stimulus onset asynchronies (SOA): ±10, ±15, ±30, ±45, ±60, ±75, ±90, ±150, ±200, ±400, ±500, ±600ms. Negative values indicated that the left hand was stimulated first, positive values indicated that the right hand was stimulated first. Within each experimental block, the presented SOA for a given trial was selected according to the participant’s performance in all the previous trials, using the adaptive PSI method [34]. Based on a Bayesian framework, this adaptive procedure estimates the posterior distribution of the parameters of interest by minimizing their expected entropy (i.e. uncertainty) trial by trial, so that the SOA selected at each trial gives the most information to estimate the parameters of interest without probing extensively all the possible SOA [23]. Two of the 4 blocks were performed with the hands in an uncrossed posture and the two other blocks in a crossed posture (i.e. arms crossed over the body midline). Within each posture condition, participants performed one block in which they had to respond according to an anatomical instruction (“*Which hand was stimulated first?*”) and one block in which a spatial instruction was used (“*From which side of space came the first stimulus?*”). The order of the *posture* and *instruction* conditions was counterbalanced across participants. Participants had to verbally report either on which hand they felt the first stimulus or from which side of space came the first stimulus of the pair, by saying “*left*” or “*right*” out loud. Participants’ responses were encoded by the experimenter on a keyboard triggering the next trial 2000 ms later. Time interval between two trials varied from 5 to 10 seconds and included the time needed to displace the laser beam on the two participant’s hands for the next trial. The task was unspeeded but the participants were instructed to be as accurate as possible. No feedback was available regarding their performance in the task.

A practice session preceded the experiment and consisted of 4 blocks of 5 trials each, one block per hand posture and per instruction condition (i.e. uncrossed vs. crossed, and “which hand” vs. “which side of space”). Only two among the largest SOA were presented during this practice session (±150 and ±200 ms). One block lasted between 10 and 15 minutes. A ten minutes break was imposed to the participants between the experimental blocks to avoid skin overheating or habituation. The whole experiment lasted two to three hours, including the threshold measurement, the training session and the experiment per se.

### 2.4. Measures

Aδ-fiber activation thresholds and stimulation intensities (corresponding to the averaged intensity used for each hand across the experimental blocks, i.e. approximately 5°C added to the Aδ-fiber activation thresholds) were measured in degrees Celcius (°C). Regarding TOJ performances, the proportion of left stimuli perceived as being presented first was computed as a function of SOA for each experimental condition. To allow comparisons between the different conditions, responses during the crossed hands posture with spatial instruction were recoded according to the stimulated hands. For each participant, data were fitted online with the logistic function, i.e. *f(x)* = 1/(1+exp(−β(x−α))), from which the parameters of interest were derived [23]. These parameters computed by the logarithm were the threshold (α) and the slope (β) of the function. In the present experiments, we were especially interested in the β parameter, which described the noisiness of the participants’ responses, i.e. the precision of their responses during the experiment [34]. The slope is indeed classically used to measure and index the impact of posture on TOJ performances both in sighted and blind participants tasks [7; 9; 18; 20–22; 30; 44–46; 54], usually by means of the just noticeable difference (JND) that denotes the SOA needed for the participants to correctly perceive the order of the two stimuli in a certain percentage of trials [30]. The α was the threshold of the function and referred to the point of subjective simultaneity (PSS), defining the SOA at which the participants reported the two stimuli as occurring first equally often (in milliseconds). Although this parameter was not relevant for our research questions, it was taken into consideration in order to investigate the presence of potential biases toward the perception of the stimuli applied to one of the hands, which could have influenced the estimation of the slope. Indeed, in the frame of the adaptive PSI method, the measures of the threshold and the slope are not completely independent and a large threshold value, indicating a bias in the perception of nociceptive stimuli towards one hand, could reduce the noisiness of the participant’s responses because of the predictability of these responses. The last update computed by the logarithm of the adaptive procedure during the task constituted the parameter estimates [35]. Because the PSI method was based on a bayesian approach, a prior probability distribution needed to be postulated, based on previous knowledge regarding the values of the parameter of interest [23]. For the present experiments, the prior distributions were set at 0 ±20 and 0.06±0.6 for the α and the β parameters, respectively [23]. Since we used an adaptive method, a third parameter was derived from the data: the mode of the presented SOA. This corresponded to the value of the SOA, among all the possible SOA, that was the most frequently presented to each participant during the adaptive procedure (in milliseconds). This additional parameter can be considered as a measure of the participant’s adaptation during the task, following the idea that smaller was the mode, better was the performance, and larger it was, worse was the performance. Indeed, this indicated that larger SOA were needed for the participant to be able to correctly discriminate between both stimuli during the task.

### 2.5. Analyses

Data were excluded from further statistical analyses if the slope of the psychometric function could not be reliably estimated during the 40 trials within one condition. Analyses were first performed on the Aδ-fibers activation threshold and stimuli intensity values, to ensure that no difference between hands or groups regarding these factors could have influenced the results. In Experiment 1, comparison of activation thresholds and stimulation intensities was made using paired t-tests with *hand* as the factor (left vs. right). In Experiment 2, we used an analysis of variance (ANOVA) for repeated measures with adding the group as second factor (sighted vs. blind).

Regarding TOJ values, one-sample t-tests were first performed to compare the PSS values to 0 for each condition of the *posture* and *instruction* factors, and each of the three groups, in order to examine the presence of potential biases. Next, the effects of the different factors on the PSS, slope and mode of the SOAs values were compared by means of analyses of variance (ANOVA) for repeated measures. For Experiment 1, *posture* (uncrossed vs. crossed) and *instruction* (anatomical vs. spatial) were used as within-participants factors. For Experiment 2, *group* (early blind vs. normally sighted) was added as a between-participants factor. Greenhouse-Geisser corrections and contrast analyses were used if necessary. Effect sizes were measured using partial Eta squared for ANOVA and Cohen’s d for t-tests. Significance level was set at *p*≤ .050.

## 3. Results

### 3.1. Experiment 1

#### 3.1.1. Threshold and intensity values

Paired sample t-tests revealed no difference between the Aδ-fiber activation threshold values of the left (M=47.80±1.52°C) vs. the right hand (M=47.40±1.77°C) in the sighted participants in Experiment 1 (t(14)=0.72, *p*=.442, d=.26). Similarly, no difference were found regarding the mean intensity of nociceptive stimuli between the left (M=52.55±1.98°C) and the right (M=52.34±1.98°C) hands (t(14)=0.39, *p*=.702, d=0.26).

#### 3.1.2. TOJ values

Results of Experiment 1 are illustrated in Figure 2. The t-tests showed that none of the PSS values from each posture and instruction conditions was significantly different from 0 (all t(14)≤.977, *p*≥.345, d≤.25). The ANOVA performed on the PSS values revealed neither a significant main effect of the posture (F(1,14)=.01, *p*=.941, ŋ^2^_*p*_<.01), nor of the instruction (F(1,14)=.56, *p*=.468, ŋ^2^_*p*_=.04), nor significant interaction between the two factors (F(1,14)<.01, *p*=.970, ŋ^2^_*p*_<.01).

**Figure 2.**
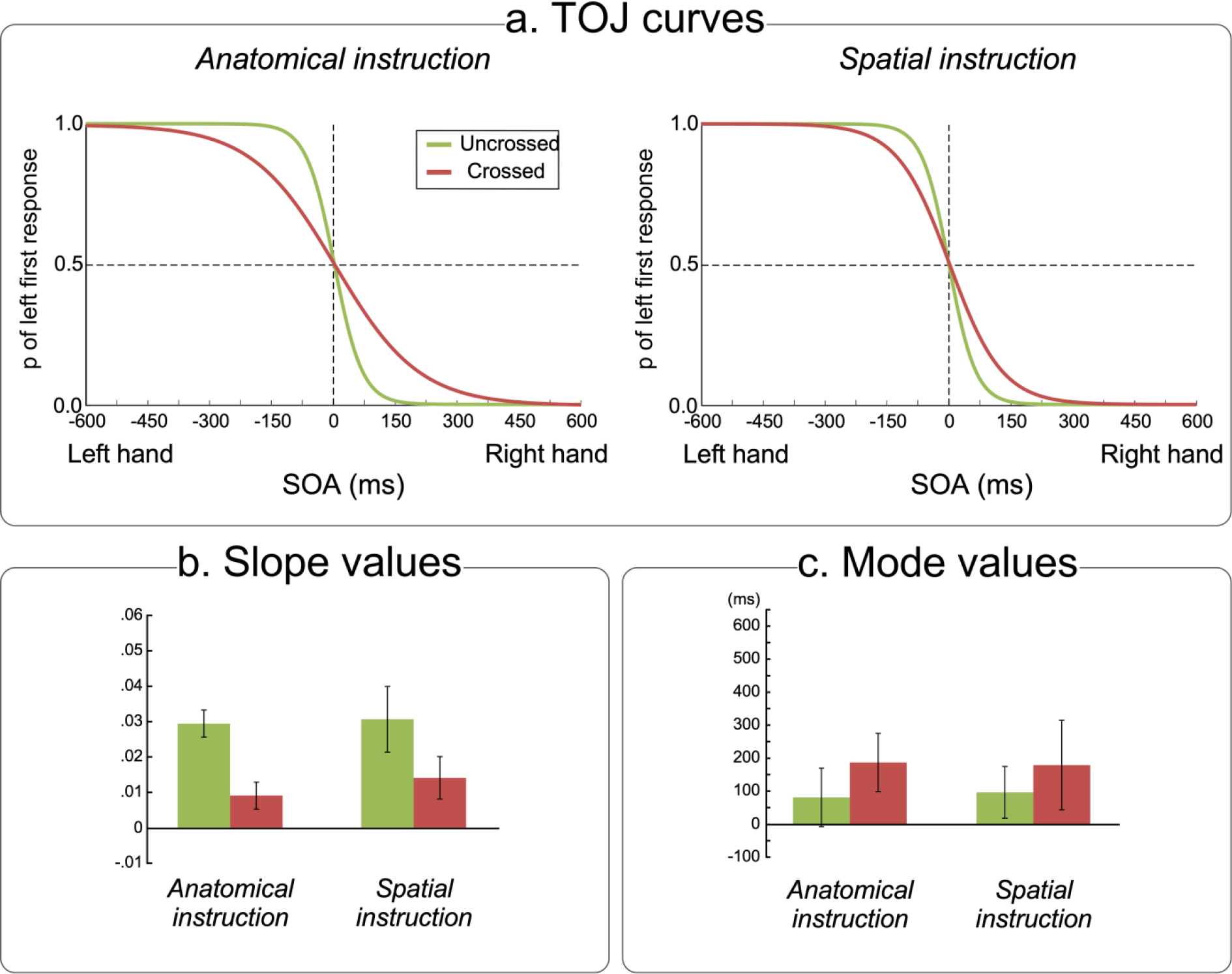
Nociceptive TOJ tasks of Experiment 1. (a) Fitted curves of the psychometric functions from the data of 15 normally sighted participants according to the posture (i.e. uncrossed vs. crossed) and according to the instruction (i.e. anatomical vs. spatial) conditions. The x-axis represents the different possible SOAs. A negative value indicates that the left hand was stimulated first and a positive value indicates that the right hand was stimulated first. The y-axis refers to the proportion of trials in which the nociceptive stimulus applied on the left hand was perceived as being presented first. The lines represent the fitted curves computed by the adaptive logarithm for the uncrossed (green) and crossed (red) conditions respectively. (b) Averaged slope values for each posture and instruction condition. The mean of the slope values was significantly lower in the crossed as compared to the uncrossed condition, whatever the instruction condition. (c) Averaged mode values of the presented SOA according to each posture and instruction condition. Error bars represent confidence intervals calculated according to Cousineau’s method for within-subject designs [17].

The analysis of the slope values showed a significant main effect of the posture (F(1,14)=37.34, *p*<.001, ŋ^2^_*p*_=.73) with no significant effect of the instruction (F(1,14=1.54, *p*=.235, ŋ^2^_*p*_=.10) and no significant interaction between both factors (F(1,14)=.33, *p*=.577, ŋ^2^_*p*_=.02). This indicated that the slope values in the crossed posture (M=.01±.01) was significantly lower than those in the uncrossed posture (M=.03±.01), whatever the instruction.

The analysis of the mode values did not reveal any significant effect of the posture (F(1,14)=2.01, *p*=.178, ŋ^2^_*p*_=.13), of the instruction (F(1,14)=.01, *p*=.921, ŋ^2^_*p*_<.01), nor any significant interaction between the two factors (F(1,14)=.08, *p*=.781, ŋ^2^_*p*_<.01).

### 3.2. Experiment 2

#### 3.2.1. Threshold and intensity values

The ANOVA performed on the Aδ-fibers activation thresholds did not show significant effect of the hand (F(1,18)=.74, *p*=.400, ŋ^2^_*p*_=.04), of the group (*F*(1,18)=0.33, *p*=.574, ŋ^2^_*p*_=.02), nor any significant interaction between the two factors (F(1,18)=.24, *p*=.628, ŋ^2^_*p*_=.01). Early blind participants thus had a similar Aδ-fibers activation threshold (M=49.75±3.91°C) than the sighted participants (M=50.55±2.06°C). Similar results were obtained regarding the stimulation intensities since the analyses showed neither significant effect of the hand (F(1,18)=2.04, *p*=.171, ŋ^2^_*p*_=.10), nor significant effect of the group (F(1,18)=.85, *p*=.369, ŋ^2^_*p*_=.01), or significant interaction between the two factors (F(1,18)=.17, *p*=.688, ŋ^2^_*p*_=.01). The averaged stimulation intensity used for the early blind group (M=54.77±2.77°C) was comparable to that for the sighted group (M=55.73±1.79°C).

#### 3.2.2. TOJ values

Results of Experiment 2 are illustrated in Figure 3 and 4. None of the PSS values were significantly different from 0 in the early blind group (all t(9)≤−.54, all *p*≥.224, d≤.17). In the sighted group, most of the PSS values were not significantly different from 0 (all t≤−1.91, *p*≥.088, d≤.60), except for the PSS value in the uncrossed posture with anatomical instruction (t(9)=−7.02, *p*<.001, d=2.22). With an averaged value of −10.12 ms (SD=4.56 ms), this indicated that their judgments were slightly biased towards the right hand in this condition. The ANOVA did not reveal significant main effect neither of the posture (F(1,18)=2.22, *p*=.154, ŋ^2^_*p*_=.11) nor of the instruction (F(1,18)=1.96, *p*=.179, ŋ^2^_*p*_=.10). In contrast, this analysis showed a significant effect of the group (F(1,18)=6.25, *p*=.022, ŋ^2^_*p*_=.26), indicating that the PSS values of normally sighted participant were more negative (M=−5.60±7.60ms) that those of the early blind participants (M=1.68±10.32ms). In addition, a significant interaction between the *posture* and the *instruction* factors was observed (F(1,18)=5.60, *p*=.029, ŋ^2^_*p*_=.24). Contrast analyses showed that during the uncrossed posture, the PSS value in the anatomical instruction condition (M=−6.05±9.57) was larger than that of the spatial instruction condition (M=−0.80±10.70ms; t(19)=−2.15, *p*=.045, d=.48). Such difference was not significant during the crossed posture (t(19)=.54, p=.593, d=.12). None of the other possible interactions was significant (all F(1,18)≤.22, all *p*≥.643, ŋ^2^_*p*_≤.01).

**Figure 3.**
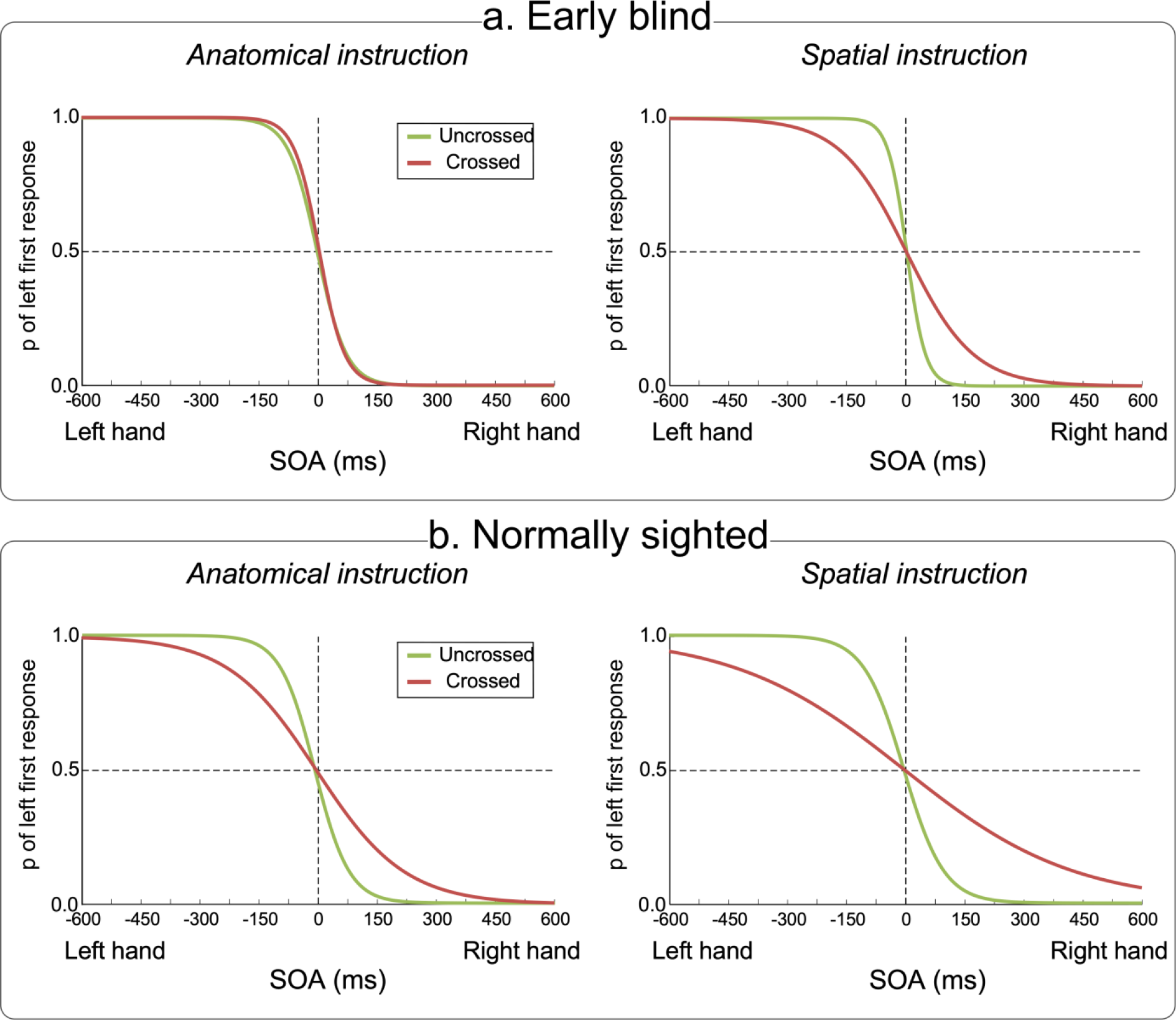
Nociceptive TOJ tasks of Experiment 2. The figure illustrates the fitted curves of the psychometric functions from data of 10 early blind (a.) and 10 sighted participants (b.) according to the posture (uncrossed in green vs. crossed in red) and the instruction conditions (anatomical on the left vs. spatial on the right side of the figure). The x-axis represents the different possible SOAs. A negative value indicates that the left hand was stimulated first and a positive value indicates that the right hand was stimulated first. The y-axis refers to the proportion of trials in which the nociceptive stimulus applied on the left hand was perceived as being presented first.

**Figure 4.**
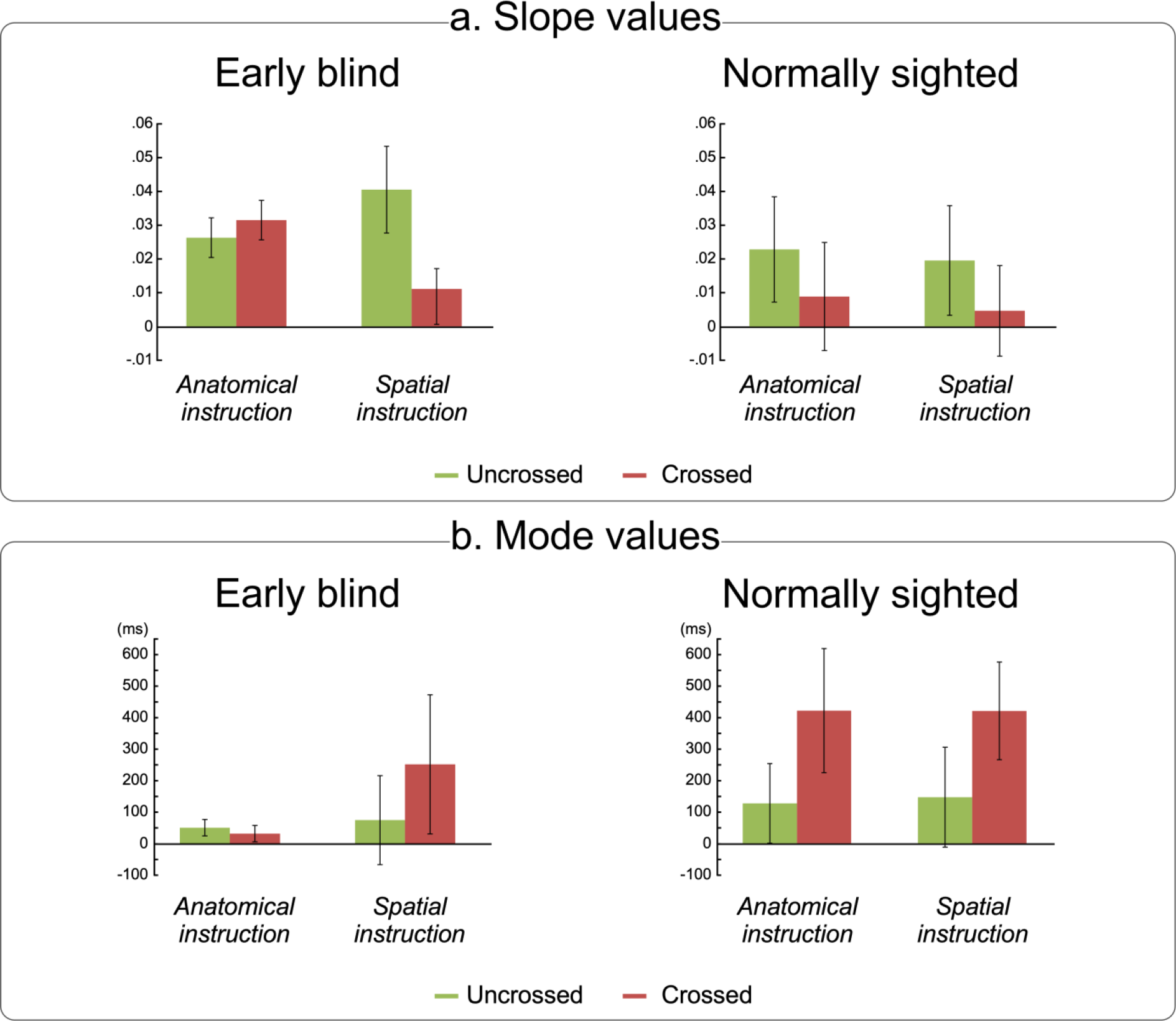
Slope and mode values of the nociceptive TOJ tasks of Experiment 2. (a) Mean slope values for each posture (uncrossed in green vs. crossed in red) and instruction conditions (anatomical on the left vs. spatial on the right part of the graphs) for respectively the early blind (left) and normally sighted participants (right). In the early blind group, a lower averaged slope value was found in the crossed as compared to the uncrossed posture condition in the spatial instruction condition, whereas the performance of this group did not significantly differ according to the posture in the anatomical instruction condition. Conversely, in the sighted group, the averaged slope value was significantly lower in the crossed as compared to the uncrossed posture condition, whatever the instruction condition. (b) Mean values of the mode of the presented SOA for each posture (uncrossed in green vs. crossed in red) and instruction conditions (anatomical on the left vs. spatial on the right part of the graphs) for respectively the early blind (left) and normally sighted participants (right). Whereas no significant difference between the conditions was evidenced in the blind group, significantly higher mode values were found in the sighted group in the crossed as compared to the uncrossed posture condition, irrespective of the instructions. Error bars represent confidence intervals calculated according to Cousineau’s method for within-subject designs [17].

Regarding the slope values, the ANOVA showed a significant main effect of the group (F(1,18)=4.80, *p*=.042, ŋ^2^_*p*_=.21), a significant main effect of the posture (F(1,18)=27.15, *p*<.001, ŋ^2^_*p*_=.60), a significant main effect of the instruction (F(1,18)=4.68, *p*=.044, ŋ^2^_*p*_=.21), and a significant interaction between *posture* and *instruction* (F(1,18)=7.23, *p*=.015, ŋ²_*p*_=.29). We also observed a significant triple interaction between all factors (F(1,18)=6.47, *p*=.020, ŋ^2^_*p*_=.26). None of the other interactions was significant (all F(1,18)≤.21, all *p*≥.653, ŋ^2^_*p*_≤.01). Next, analyses were run separately in each group of participants with *posture* and *instruction* as within factors. For the early blind participants, we observed a significant main effect of the posture (F(1,9=6.45, *p*=.032, ŋ^2^_*p*_=.42), a significant interaction between the *posture* and *instruction* factors (F(1,9)=8.27, *p*=.018, ŋ^2^_*p*_=.48), but no significant effect of the instruction (F(1,9)=2.26, *p*=.167, ŋ^2^_*p*_=.20). Contrast analyses revealed that, whereas there was no significant difference between the two postures in the anatomical instruction condition (t(9)=−1.00, *p*=.343, d=0.22; M uncrossed=.03±.01, M crossed=.03±.02), such difference was significant in the spatial instruction condition (t(9)=3.08, *p*=.013, d=0.68). Slope values were indeed smaller in the crossed posture (M=.01±.01) than in the uncrossed posture conditions (M=.04±.03) in the early blind group. In contrast, in the normally sighted group, only a significant main effect of the posture was observed (F(1,9)=64.82, *p*<.001, ŋ^2^_*p*_=.88) with neither significant effect of the instruction (F(1,9)=2.44, p=.150, ŋ^2^_*p*_=.21) nor significant interaction between the two factors (F(1,9)=.03, *p*=.87, ŋ^2^_*p*_<.01). For the normally sighted participants, the slope values in the crossed posture condition (M=0.01±.00) were significantly lower than those in the uncrossed posture condition (M=0.02±.01). To summarize, results showed that the posture affected performances of the normally sighted participants whatever the instruction condition, whereas early blind participants’ performance was only affected by the crossed posture when a spatial response was required.

Finally, the ANOVA performed on the mode of the presented SOA values showed a significant main effect of the posture (F(1,18)=34.3, *p*<.001, ŋ^2^_*p*_=.66), a significant main effect of the group (F(1,18)=14.26, *p*=.001, ŋ^2^_*p*_=.44) and a significant interaction between these two factors (F(1,18)=10.95, *p*=.004, ŋ^2^_*p*_=.38). None of the other comparisons was significant (all F(1,18)≤1.46, all *p*≥.243, all ŋ^2^_*p*_≤.08). Analyses were then run separately in each group. In the early blind group, the ANOVA showed neither significant effect of the posture (F(1,9)=3.13, *p*=.11, ŋ^2^_*p*_=.26), nor of the instruction (F(1,9)=3.19, *p*=.11, ŋ^2^_*p*_=.26). The interaction between these two factors was just a bit above significance level (F(1,9)=4.01, *p*=.070, ŋ^2^_*p*_=.31). In the sighted group, a significant effect of the posture was observed (F(1,9)=43.67, *p*<.001, ŋ^2^_*p*_=.83), the mode value in crossed posture being higher (M=420.50±137.24) than that in the uncrossed posture condition (M=136.25±122.24). The effect of the instruction did not reach significance (F(1,9)=.01, *p*=.920, ŋ^2^_*p*_<.01), neither the interaction between the two factors (F(1,9)=.02, *p*=.900, ŋ^2^_*p*_<.01).

## 4. Discussion

The objectives of the present experiments were to investigate the role of visual experience and cognitive goals in shaping the spatial representations of nociceptive stimuli. To this aim, we compared performances of normally sighted and early blind participants during temporal order judgments of nociceptive stimuli, during which instructions favoured to use either an anatomical (“*which hand?*”) or a spatial (“*which space?*”) representation. Importantly, TOJ tasks were performed with the hands either uncrossed or crossed over body midline, the latter condition being intended to generate a mismatch between somatotopic and spatiotopic representations. Results showed that sighted participants’ performances were decreased in the crossed hands posture independently of which reference frame was task-relevant. On the contrary, performances of early blind participants were only affected by crossing the hands when they were requested to use a spatial response, while their performance was insensitive to the posture under anatomical instruction.

Influence of crossing the hands during cognitive tasks has been recurrently described in normally sighted people for both tactile and nociceptive stimuli [7; 9; 18; 20–22; 30; 44–46; 54]. Such an effect was interpreted as reflecting the ability of the brain to code the spatial location of somatosensory inputs according to spatiotopic reference frames [30]. Spatiotopic mapping has also been suggested to represent an important process, affording a common spatial framework for inputs from somatic and extra-somatic sensory modalities to be integrated in a peripersonal representation of the body [22; 25]. In this line, nociceptive and visual stimuli would optimize detection and reaction against physical threats around the body [38]. Together, this would suggest a default and major role of spatiotopic representations over somatotopic ones in the perception of somatosensory stimuli. However, the present data offers a new insight on this assumed dominance.

In normally sighted participants, results confirmed that nociceptive stimuli were mapped into spatiotopic representations taking body posture into account, in addition to be coded according to somatotopic representations [22; 45]. Since TOJ performance was similarly impaired under the two different task instructions, this indicated that both somatotopic and spatiotopic representations might be co-activated by default during spatial localization processing of nociceptive inputs. Hence, adapting cognitive goals by changing task instruction to stress external space-based coordinates was not enough to give advantage to the spatiotopic frame of reference and attenuate the crossing hands deficit in our experiments (see [21] for similar results with tactile stimuli). On the contrary, studies on tactile processing suggested that spatial coding of somatosensory stimuli was actually weighted depending on task demands and cognitive goals [6; 7; 9; 26]. Specifically, the results of these experiments indicated that the weight accorded to the spatiotopic representation could be attenuated under some conditions, but externally defined coordinates still had a robust influence on the participants’ responses during tactile processing. Accordingly, the present data suggest that nociceptive stimuli are automatically coded according to both somatotopic and spatiotopic reference frames, but that the weight respectively given to each reference frames during the localization processing of nociception would be more dependent on contextual factors such as cognitive goals [7; 8].

Additionally, results in early blind participants indicated that the assumed weighted activation of the spatiotopic reference frame of nociceptive stimuli might be driven by early visual experience. Indeed, a default advantage of somatotopic representations in congenitally blind people during touch localization had been recurrently suggested by means of behavioural as well as electrophysiological and neuroimaging data [19; 20; 43]. However, regarding nociception and pain, being able to consider the position of the limbs in external space is of primary importance to protect the body from potential physical threats. Then, any individual should be able to take such spatial information into account, if not automatically, at least when it is relevant for the ongoing situation. Accordingly, we observed that a crossing hands deficit emerged in early blind participants in the spatial instruction condition, confirming that early blind participants were able to activate and use spatiotopic representations. Indeed, this finding suggests that the spatial representations at play when localizing nociceptive events could be adapted according to the task requirements. We showed that making external space relevant by changing the instruction was sufficient to highlight the use of the spatiotopic reference frame in this group during TOJ tasks. These data are in line with a very recent study having shown the exact same pattern of results using a tactile TOJ task in early blind and normally sighted participants [21]. Other studies showed that, in tactile motor coordination tasks, spatial external coordinates were actually used by congenitally blinds when task demands prioritized the external space reference [18; 31]. Taken together, these studies suggest a differential balance between the activation by default of the somatotopic and spatiotopic frames of reference in normally sighted and early blind participants during somatosensory perception [18; 20; 21]. Following this idea, early blind individuals might have better abilities to inhibit the spatial responses when it is irrelevant for the ongoing activity, while formatting responses according to external space would be resistant to inhibition in sighted people, even when irrelevant to current behavioural goals.

The present studies therefore suggest that early visual experience shapes the way spatiotopic mapping of nociceptive stimuli develops, resulting in qualitatively different ways of processing nociception and pain in adulthood between early blinds and normally sighted individuals. Perhaps as a result, some quantitative differences were observed between congenitally blind and normally sighted individuals regarding the perception of pain. For instance, Slimani et al. [47; 48] observed lower thresholds for heat and cold pain, as well as faster reaction times to stimulations mediated by C-, but not Aδ-, fibers in congenitally blinds as compared to sighted controls. These authors have linked this “*hypersensitivity*” to pain in this population to higher levels of anxiety and enhanced attention to painful stimuli [33]. Altogether, the present and previous experiments on early blindness highlight that the plasticity of the nociceptive system depends on early sensory experience from any sensory modalities, including non-somatic ones.

The present data point that nociceptive and tactile modalities seem to share, at least partially, the same spatial representations. However, while the cortical substrates underlying the spatial representation of touch were extensively studied [3; 5; 12; 15; 20; 28; 39; 49; 50; 53], they are still poorly understood for nociception [38; 51]. Premotor and posterior parietal brain areas have been largely shown to be involved in the spatial coding of tactile stimuli [5; 12; 20; 39; 50; 53] and in visuo-tactile crossmodal interactions [4; 15; 28; 39; 40]. Nociceptive and painful stimuli were also shown to elicit brain activity in premotor and posterior parietal areas [see review in 2]. On the other hand, it was suggested that the cortical regions classically observed in response to nociceptive and painful stimuli, such as cingulate and operculo-insular cortices, were usually associated to a broader brain network involved in the detection of any sensory stimulus that might have an impact on the body’s integrity [e.g. 37]. Therefore, we hypothesize that premotor and posterior parietal brain areas might be associated to that network with the aim of mapping threatening objects into external spatial coordinates in order to prepare spatially guided actions to protect the body’s integrity.

The present data also offer a new insight on the so-called “*crossing hands analgesia*” according to which crossing the hands over the body midline decreases cortical responses to nociceptive stimuli and the perception of their intensity [27; 51]. Gallace et al. [27] interpreted this effect as reflecting a disruption of nociceptive processing in the brain. Accordingly, during unusual body posture such as when the hands are crossed, privileged connections with the nociceptive system would not be engaged, especially those involved in spatial perception, resulting in a decrease of the cortical responses, which would in turns impede the processing of the intensity and its perception. However, based on present results, we propose an alternative hypothesis according to which, when the different reference frames are conflicting, the less accurate TOJ performance would reflect the effort that the brain has to make to prioritize the relevant reference frame and inhibit the irrelevant spatial code. We therefore hypothesize that the so-called analgesic effect observed during the crossed hands posture results from a lack of processing resources left out by the competition between the different spatial representations. In other words, resolving the conflict between the different reference frames and selecting the relevant spatial response requests attentional resources [7] that are less available to process other stimulus features such as its intensity. Manipulating the attentional load was indeed shown to modulate nociceptive processing [36].

In conclusion, the present studies emphasize that localization of nociceptive stimuli is based on multiple mapping systems taking into account both the space of the body and external space. We showed that activating the different spatial reference frames is automatic but disentangling between the different spatial responses where there is a mismatch requires effort and depends on developmental and contextual weighting. Further experiments will be needed to disclose the cortical mechanisms underlying the spatial representations of pain and their impairments during pathological pain [24].

## 5. Acknowledgements

CV, ADV and VL are supported by the Funds for Scientific Research of the French-speaking Community of Belgium (F.R.S.-FNRS). The authors state that there is no conflict of interest.

## References

[1] Andersson JLR, Lilja, A., Hartvig, P., Langström, B., Gordh, T., Kandwerker, H., Torebjörk, E. Somatotopic organization along the central sulcus, for pain localization in humans, as revealed by positron emission tomography. Experimental Brain Research 1997;117:192–199.

[2] Apkarian AV, Bushnell MC, Treede R-D, Zubieta J-K. Human brain mechanisms of pain perception and regulation in health and disease. European Journal of Pain 2005;9(4):463–463.

[3] Avillac M, Deneve S, Olivier E, Pouget A, Duhamel J-R. Reference frames for representing visual and tactile locations in parietal cortex. Nat Neurosci 2005;8(7):941–949.

[4] Avillac M, Deneve S, Olivier E, Pouget A, Duhamel JR. Reference frames for representing visual and tactile locations in parietal cortex. Nat Neurosci 2005;8(7):941–949.

[5] Azanon E, Longo MR, Soto-Faraco S, Haggard P. The posterior parietal cortex remaps touch into external space. Curr Biol 2010;20(14):1304–1309.

[6] Badde S, Heed T. Towards explaining spatial touch perception: Weighted integration of multiple location codes. Cogn Neuropsychol 2016;33(1-2):26–47.

[7] Badde S, Heed T, Roder B. Processing load impairs coordinate integration for the localization of touch. Atten Percept Psychophys 2014;76(4):1136–1150.

[8] Badde S, Heed T, Roder B. Integration of anatomical and external response mappings explains crossing effects in tactile localization: A probabilistic modeling approach. Psychon Bull Rev 2016;23(2):387–404.

[9] Badde S, Roder B, Heed T. Flexibly weighted integration of tactile reference frames. Neuropsychologia 2015;70:367–374.

[10] Baumgartner U, Iannetti GD, Zambreanu L, Stoeter P, Treede RD, Tracey I. Multiple somatotopic representations of heat and mechanical pain in the operculo-insular cortex: a high-resolution fMRI study. J Neurophysiol 2010;104(5):2863–2872.

[11] Bingel U, Lorenz J, Glauche V, Knab R, Gläscher J, Weiller C, Büchel C. Somatotopic organization of human somatosensory cortices for pain: a single trial fMRI study. NeuroImage 2004;23(1):224–232.

[12] Bolognini N, Maravita A. Proprioceptive alignment of visual and somatosensory maps in the posterior parietal cortex. Curr Biol 2007;17(21):1890–1895.

[13] Brooks JC, Zambreanu L, Godinez A, Craig AD, Tracey I. Somatotopic organisation of the human insula to painful heat studied with high resolution functional imaging. Neuroimage 2005;27(1):201–209.

[14] Brozzoli C, Ehrsson HH, Farne A. Multisensory representation of the space near the hand: from perception to action and interindividual interactions. Neuroscientist 2014;20(2):122–135.

[15] Brozzoli C, Gentile G, Petkova VI, Ehrsson HH. fMRI Adaptation Reveals a Cortical Mechanism for the Coding of Space Near the Hand. J Neurosci 2011;31(24):9023–9031.

[16] Churyukanov M, Plaghki L, Legrain V, Mouraux A. Thermal detection thresholds of Adelta- and C-fibre afferents activated by brief CO2 laser pulses applied onto the human hairy skin. PLoS One 2012;7(4):e35817.

[17] Cousineau D. Confidence intervals in within-subject designs: a simpler solution to Loftus and Masson’s method. Tutor Quant Methods Psychol 2005;1(1):42–45.

[18] Crollen V, Albouy G, Lepore F, Collignon O. How visual experience impacts the internal and external spatial mapping of sensorimotor functions. Sci Rep 2017;7(1):1022.

[19] Crollen V, Collignon O. Embodied space in early blind individuals. Front Psychol 2012;3:272.

[20] Crollen V, Lazzouni L, Rezk M, Bellemare A, Lepore F, Collignon O. Visual Experience Shapes the Neural Networks Remapping Touch into External Space. J Neurosci 2017;37(42):10097–10103.

[21] Crollen V, Spruyt S, Mahau P, Bottini R, Collignon C. How visual experience and task context modulate the use of internal and external spatial coordinate for perception and action. JEP-HPP In press.

[22] De Paepe AL, Crombez G, Legrain V. From a somatotopic to a spatiotopic frame of reference for the localization of nociceptive stimuli. PLoS One 2015;10(8):e0137120.

[23] Filbrich L, Alamia A, Burns S, Legrain V. Orienting attention in visual space by nociceptive stimuli: investigation with a temporal order judgment task based on the adaptive PSI method. Exp Brain Res 2017;235:2069–2079.

[24] Filbrich L, Alamia A, Verfaille C, Berquin A, Barbier O, Libouton X, Fraselle V, Mouraux D, Legrain V. Biased visuospatial perception in complex regional pain syndrome. Scientific Reports 2017;7(1):9712.

[25] Filbrich L, Halicka M, Alamia A, Legrain V. Investigating the spatial characteristics of the crossmodal interaction between nociception and vision using gaze direction. Conscious Cogn 2018;57:106–115.

[26] Gallace A, Soto-Faraco S, Dalton P, Kreukniet B, Spence C. Response requirements modulate tactile spatial congruency effects. Exp Brain Res 2008;191(2):171–186.

[27] Gallace A, Torta DM, Moseley GL, Iannetti GD. The analgesic effect of crossing the arms. Pain 2011;152(6):1418–1423.

[28] Graziano MSA, Hu XT, Gross CG. Visuospatial Properties of Ventral Premotor Cortex. J Neurophysiol 1997;77(5):2268–2292.

[29] Haggard P, Iannetti GD, Longo MR. Spatial sensory organization and body representation in pain perception. Curr Biol 2013;23(4):R164–176.

[30] Heed T, Azañon E. Using time to investigate space: a review of tactile temporal order judgments as a window onto spatial processing in touch. Frontiers in Psychology 2014;5:76.

[31] Heed T, Roder B. Motor coordination uses external spatial coordinates independent of developmental vision. Cognition 2014;132(1):1–15.

[32] Henderson LA, Gandevia SC, Macefield VG. Somatotopic organization of the processing of muscle and cutaneous pain in the left and right insula cortex: a single-trial fMRI study. Pain 2007;128(1-2):20–30.

[33] Holten-Rossing S, Slimani H, Ptito M, Danti S, Kupers R. Uncertainty about the intensity of impending pain increases ensuing pain responses in congenital blindness. Behav Brain Res 2018;346:41–46.

[34] Kingdom FAA, Prins N. Psychophysics - A practical introduction. London: Elsevier Academic Press, 2010.

[35] Kontsevich LL, Tyler CW. Bayesian adaptive estimation of psychometric slope and threshold. Vision Res 1999;39:2729–2737.

[36] Legrain V, Crombez G, Plaghki L, Mouraux A. Shielding cognition from nociception with working memory. Cortex 2013;49(7):1922–1934.

[37] Legrain V, Iannetti GD, Plaghki L, Mouraux A. The pain matrix reloaded: a salience detection system for the body. Prog Neurobiol 2011;93(1):111–124.

[38] Legrain V, Torta DM. Cognitive psychology and neuropsychology of nociception and pain. In: G Pickering, S Gibson, editors. Pain, Emotion and Cognition: A complex Nexus. Paris: Springer 2015. pp. 2–20.

[39] Lloyd DM, Shore DI, Spence C, Calvert GA. Multisensory representation of limb position in human premotor cortex. Nat Neurosci 2003;6(1):17–18.

[40] Makin TR, Holmes NP, Zohary E. Is that near my hand? Multisensory representation of peripersonal space in human intraparietal sulcus. J Neurosci 2007;27(4):731–740.

[41] Nicholls ME, Thomas NA, Loetscher T, Grimshaw GM. The Flinders Handedness survey (FLANDERS): a brief measure of skilled hand preference. Cortex 2013;49(10):2914–2926.

[42] Penfield W, Rasmussen T. The cerebral cortex of man; a clinical study of localization of function. Oxford, England: Macmillan, 1950.

[43] Roder B, Focker J, Hotting K, Spence C. Spatial coordinate systems for tactile spatial attention depend on developmental vision: evidence from event-related potentials in sighted and congenitally blind adult humans. Eur J Neurosci 2008;28(3):475–483.

[44] Röder B, Rösler F, Spence C. Early Vision Impairs Tactile Perception in the Blind. Current Biology 2004;14(2):121–124.

[45] Sambo CF, Torta DM, Gallace A, Liang M, Moseley GL, Iannetti GD. The temporal order judgement of tactile and nociceptive stimuli is impaired by crossing the hands over the body midline. Pain 2013;154(2):242–247.

[46] Shore DI, Spry E, Spence C. Confusing the mind by crossing the hands. Cogn Brain Res 2002;14:153–163.

[47] Slimani H, Danti S, Ricciardi E, Pietrini P, Ptito M, Kupers R. Hypersensitivity to pain in congenital blindness. Pain 2013;154(10):1973–1978.

[48] Slimani H, Plaghki L, Ptito M, Kupers R. Pain hypersensitivity in congenital blindness is associated with faster central processing of C-fibre input. Eur J Pain 2016;20(9):1519–1529.

[49] Soto-Faraco S, Azanon E. Electrophysiological correlates of tactile remapping. Neuropsychologia 2013;51(8):1584–1594.

[50] Takahashi T, Kansaku K, Wada M, Shibuya S, Kitazawa S. Neural correlates of tactile temporal-order judgment in humans: an fMRI study. Cereb Cortex 2013;23(8):1952–1964.

[51] Torta DM, Diano M, Costa T, Gallace A, Duca S, Geminiani GC, Cauda F. Crossing the line of pain: FMRI correlates of crossed-hands analgesia. J Pain 2013;14(9):957–965.

[52] Vallar G, Maravita A. Personal and extrapersonal spatial perception. In: GG Berntson, JT Cacioppo, editors. Handbook of Neuroscience for the Behavioral Sciences, Vol. 1. New York: John Wiley & Sons, 2009. pp. 322–336.

[53] Wada M, Takano K, Ikegami S, Ora H, Spence C, Kansaku K. Spatio-temporal updating in the left posterior parietal cortex. PLoS One 2012;7(6):e39800.

[54] Yamamoto S, Kitazawa S. Reversal of subjective temporal order due to arm crossing. Nature Neuroscience 2001;4(7):759–765.

